# Regulatory Sharing Between Estrogen Receptor α Bound Enhancers

**DOI:** 10.1101/2020.03.18.997403

**Authors:** Julia B. Carleton, Matthew Ginley-Hidinger, Kristofer C. Berrett, Ryan M. Layer, Aaron R. Quinlan, Jason Gertz

## Abstract

The human genome encodes an order of magnitude more gene expression enhancers than promoters, suggesting that most genes are regulated by the combined action of multiple enhancers. We have previously shown that neighboring estrogen-responsive enhancers, which are approximately 5,000 basepairs apart, exhibit complex synergistic contributions to the production of an estrogenic transcriptional response. Here we sought to determine the molecular underpinnings of the observed enhancer cooperativity. We generated genetic deletions of individual estrogen receptor α (ER) bound enhancers and found that enhancers containing full estrogen response element (ERE) motifs control ER binding at neighboring sites, while enhancers with pre-existing histone acetylation/accessibility confer a permissible chromatin environment to the neighboring enhancers. Genome engineering revealed that a cluster of two enhancers with half EREs could not compensate for the lack of a full ERE site within the cluster. In contrast, two enhancers with full EREs produced a transcriptional response greater than the wild-type locus. By swapping genomic sequences between enhancers, we found that the genomic location in which a full ERE resides strongly influences enhancer activity. Our results lead to a model in which a full ERE is required for ER recruitment, but the presence of a pre-existing active chromatin environment within an enhancer cluster is also needed in order for estrogen-driven gene regulation to occur.

## Introduction

Regulation of gene expression is a fundamental task underlying biological processes such as development and disease progression. Promoter-distal gene regulatory enhancers play a central role in metazoan gene regulation and contain binding sites for transcription factors (TFs) that recruit cofactors and influence gene expression. Most genes in the human genome are likely regulated by multiple enhancers (Long et al., 2016; Spitz and Furlong, 2012). For example, the ENCODE consortium found that an average of 3.9 distal elements are involved in long-range interactions with each transcription start site (Consortium, 2012). While multiple enhancers often combine to regulate gene expression, the molecular details of how these enhancers work together remains poorly understood, partially due to a paucity of functional studies. Understanding how multiple enhancers molecularly communicate with each other and their target gene promoter represents a major open question in gene regulation.

The most commonly observed model for how enhancers combine to regulate gene expression has them acting in an independent or additive manner, allowing for elements to evolve independently, which can lead to divergence in tissue-specific expression patterns and gene expression levels. As an example, multiple independent enhancers regulate the β-globin locus in mice (Bender et al., 2012). Enhancers can also act in a synergistic or cooperative manner to influence gene expression (Yuh and Davidson, 1996). Leddin et al. found that in order for PU.1 to bind at one of its upstream enhancers in myeloid cells and auto-regulate expression, a second enhancer must be active. This second enhancer likely maintains accessible chromatin at the neighboring enhancer, enabling PU.1 to bind (Leddin et al., 2011). Enhancers can also work together to maintain a favorable 3D chromatin architecture and promote transcription factor recruitment, as observed at the *Igk* locus in B cells (Fulton and van Ness, 1994; Proudhon et al., 2016). However, the molecular details behind enhancer interactions and features that dictate independence and cooperativity remain relatively unknown.

Estrogen signaling through estrogen receptor α (ER) is a relevant model system to study combinatorial gene regulation. ER binds the genome in an estrogen-dependent manner, with the majority of binding occurring distally from promoters (Gertz et al., 2012). The majority of genes upregulated upon estrogen treatment have multiple ER binding sites nearby (Carleton et al., 2017), indicating that multiple sites might be required for coordinating the transcriptional response to estrogen. We previously developed a CRISPR interference based method, termed enhancer interference (Carleton et al., 2018), to study enhancer relationships and identified two types of collaborative enhancer relationships: 1) hierarchical, where one predominant site contributes the majority of the estrogen response and another supportive site can contribute only when the predominant site is active, and 2) synergy, where a pair of sites is completely necessary for the estrogen response and neither site can contribute in isolation (Carleton et al., 2017). Paradoxically, when the same ER binding sites are targeted by CRISPRa fusions in the absence of estrogens, the enhancers work independently to regulate gene expression (Ginley-Hidinger et al., 2019). Taken together these findings lead to a model in which enhancers are cooperating in *cis*, positively impacting one another when activated by ER, but communicate independently with the target gene promoter when directly bound by a synthetic transcriptional activator. To resolve this apparent contradiction and advance our understanding of enhancer synergy, it is important to determine how these enhancers are molecularly cooperating when cells are treated with estrogens.

In this study, we explore how neighboring ER-bound enhancers impact one another. By functionally dissecting these relationships at 2 estrogen-responsive genes, we discovered regulatory sharing between ER-bound enhancers. Deletion of ER-bound enhancers decreased ER binding, histone acetylation, and chromatin accessibility at neighboring enhancers. Through the use of genome engineering approaches, we also investigated the role of sequence and genomic location in determining the contributions of these enhancers to estrogen-induced gene expression. We find that the presence of a full estrogen response element (ERE), ER’s preferred DNA binding sequence, in at least one enhancer is required for these genes’ transcriptional responses to estrogen. The location of the ERE containing enhancer region is also important; placement of an ERE within a region harboring histone acetylation prior to estrogen induction greatly increases the transcriptional response. Overall, we discovered that these ER-bound enhancers are cooperating at a molecular level to combine ER recruitment and permissible chromatin and drive the estrogen transcriptional response.

## Results

### Collaborative regulation by neighboring ER-bound enhancers may be a pervasive feature of the estrogen transcriptional response

*MMP17*, *FHL2*, and *CISH* each harbor pairs of ER binding sites (ERBS) that work together synergistically (Carleton et al., 2017). These ERBS pairs are within 5 kilobasepairs (kb) of each other, which led us to examine whether neighboring ERBS are a common feature of estrogen-regulated genes. To determine whether ER binding sites tend to cluster in the genome, we examined the distance between ERBS pairs using the set of 8,621 ERBS bound in Ishikawa, a human endometrial cancer cell line, by ChIP-seq (Gertz et al., 2013). We found that 42% of ERBS have a neighboring site within 10 kb (Figure 1A). In order to determine if this clustering of ERBS is a unique feature, we analyzed clustering of ERBS compared to controls of randomly subsampled DNaseI hypersensitive sites or CTCF binding sites in Ishikawa cells (8,621 loci). We assigned each individual site to a window that contained 10 kb upstream and downstream of the site and sites within 10 kb of each other were merged into a cluster. We found that only 5-8% of windows for randomly sampled CTCF bound sites and DNaseI hypersensitive sites contained more than one site; however, 22% of windows had more than one ERBS (Figure 1B), indicating a substantial enrichment for clustering of ERBS compared to other regulatory regions. ERBS clustering has been previously observed in MCF-7 cells (Bojcsuk et al., 2017; Saravanan et al., 2020), a human breast cancer cell line, and our results indicate that ERBS cluster in endometrial cancer cells as well.

**Figure 1.**
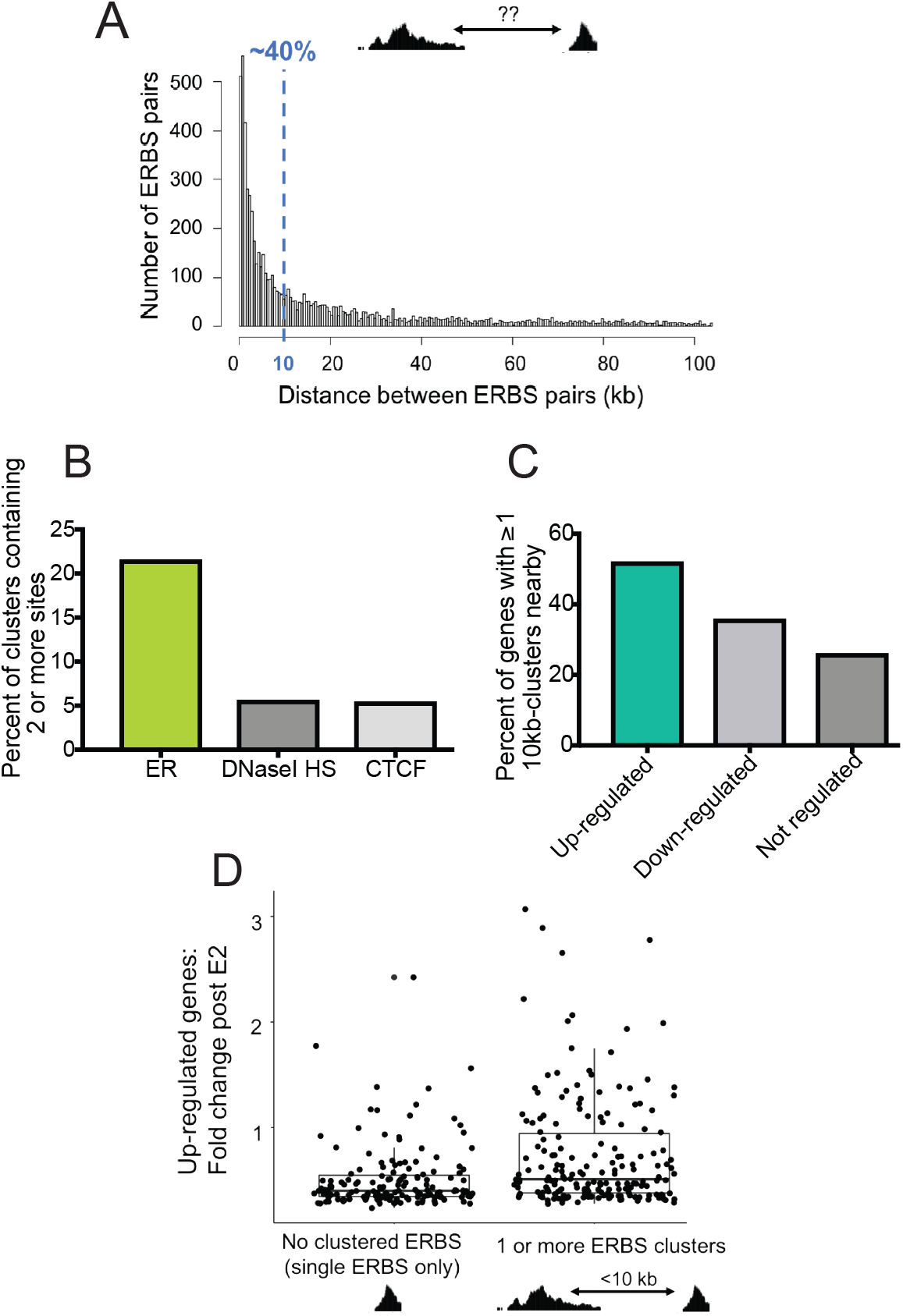
Clusters of ERBS are enriched near up-regulated genes. **A)** Histogram depicts distance between an ERBS and its closest neighboring site for all 8,621 sites bound in Ishikawa cells. **B)** ERBS, CTCF sites bound in Ishikawa, and DNaseI HS sites were each merged into 10-kb windows throughout the genome and the number of windows containing multiple sites for each feature is shown. **C)** The percent of upregulated, downregulated, and not regulated genes that have at least one cluster containing multiple ERBS is shown. **D)** Boxplot shows the relationship between fold change in response to estrogen for up-regulated genes containing either dispersed sites (0 clusters) or at least 1 cluster containing multiple ERBS within 100 kb of the TSS.

In order to better understand the relationship between clustered ERBS and gene expression, we examined where clusters are located in the genome relative to genes up-, down-, or not-regulated by estrogen as defined by RNA-seq (Gertz et al., 2013). We found that 52% of up-regulated genes have a cluster of 2 or more ERBS nearby, compared to only 35% of down-regulated genes and 26% of not-regulated genes (Figure 1C). We then compared the fold change in 17β-estradiol (E2) response for up-regulated genes with no clusters to genes with at least 1 cluster within 100 kb of the transcription start site (TSS) and found that genes with clustered ERBS nearby tended to have greater fold changes in response to E2 (Figure 1D) (p-value = 3.93 × 10^−6^, Wilcoxon signed-rank test). These results suggest that ERBS clusters may be more effective at producing gene expression responses to estrogen compared to solitary ERBS.

### Neighboring ER-bound enhancers of CISH impact each other’s ER occupancy and chromatin environment

To determine how neighboring enhancers collaborate to produce an estrogen response, we focused on two ERBS near estrogen-responsive genes *CISH* and *MMP17* (Figure 2), which have previously been shown to be required for the transcriptional response to E2 in Ishikawa cells (Carleton et al., 2017). Both genes contain a similar structure where one site in the pair has a full canonical estrogen response element (ERE) motif whereas the other site has a half ERE; however, the regulatory logic of each pair is a different type of non-independence (Figure 2A-B). At *CISH*, sites interact synergistically and both ERBS are equally necessary for the estrogen transcriptional response. At *MMP17*, a hierarchical relationship exists between ERBS, where one predominant site contributes the majority of the response and the other supportive site can contribute only when the predominant site is active (Carleton et al., 2017). The ER-bound sites display differential binding of additional transcription factors (Figure 2C-D), which may in part shape their relationships to gene expression and to each other. To determine how each ERBS contributes to its respective pair, we generated two independent cell lines for each region of interest, with each line containing a deletion of one of the four ERBS. Deletions were generated using guide RNAs flanking the ER ChIP-seq peak, which removed 125-225 basepairs (bp) of sequence.

**Figure 2.**
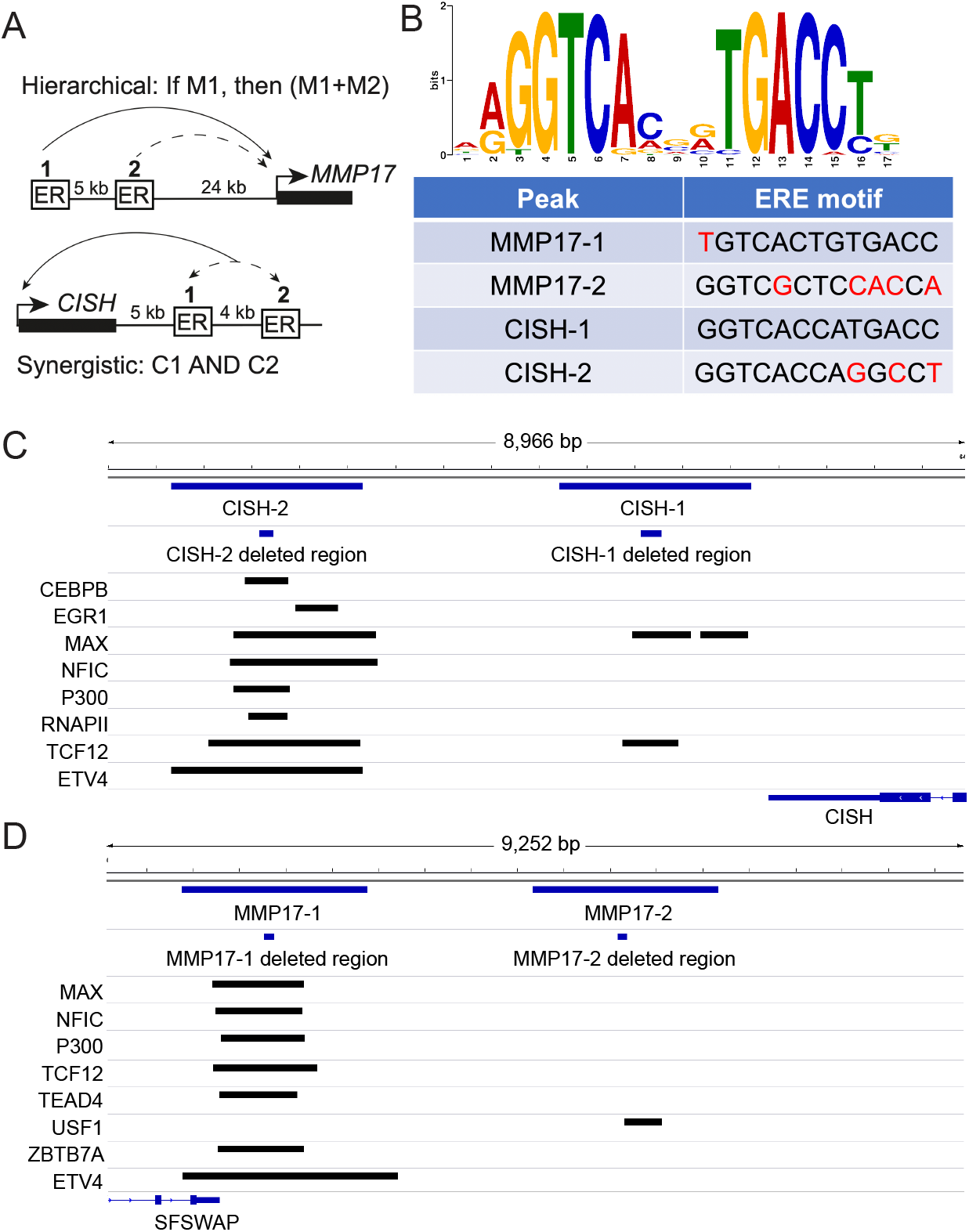
Features of enhancers controlling the estrogen transcriptional responses of *MMP17* and *CISH*. Hierarchical enhancers are upstream of *MMP17*(A). MMP17-1 contains a strong ERE (B, red letters indicate mismatches) and acts as the predominant site, while MMP17-2 has a half site and can contribute only when MMP17-1 is active. Synergistic enhancers are downstream of *CISH*(A). CISH-1 contains a canonical ERE motif (B) while CISH-2 does not. Both sites are equally necessary for the transcriptional response of *CISH*. C-D) Browser tracks show binding of additional factors to *MMP17* and *CISH* ER-bound enhancers as well as the locations of the deleted regions.

Since previous work has found that an enhancer can affect chromatin and TF binding at a neighboring enhancer, we hypothesized that ER-bound enhancers may affect each other via similar mechanisms. We performed ATAC-seq and ChIP-seq for ER and H3K27ac in the enhancer deletion lines after treatment with E2 or vehicle (DMSO). As expected, deletion of a central 225 bp of CISH-1 led to a complete loss of ER binding and large reductions in accessibility and H3K27ac at CISH-1 compared to wild-type levels (Figure 3A-D). Loss of CISH-1 also led to a 90% reduction in ER binding at CISH-2 (Figure 3B); however, H3K27ac at CISH-2 was not significantly affected either with or without E2 treatment (Figure 3C). Similar to H3K27ac, CISH-1 deletion reduced accessibility of CISH-2 by approximately 33% (p-value > 0.05) regardless of treatment (Figure 3D). These results suggest that CISH-1 participates in the synergistic relationship between CISH-1 and CISH-2 by controlling ER binding to the locus, but minimally contributes to accessible and active chromatin of the enhancer neighborhood.

**Figure 3.**
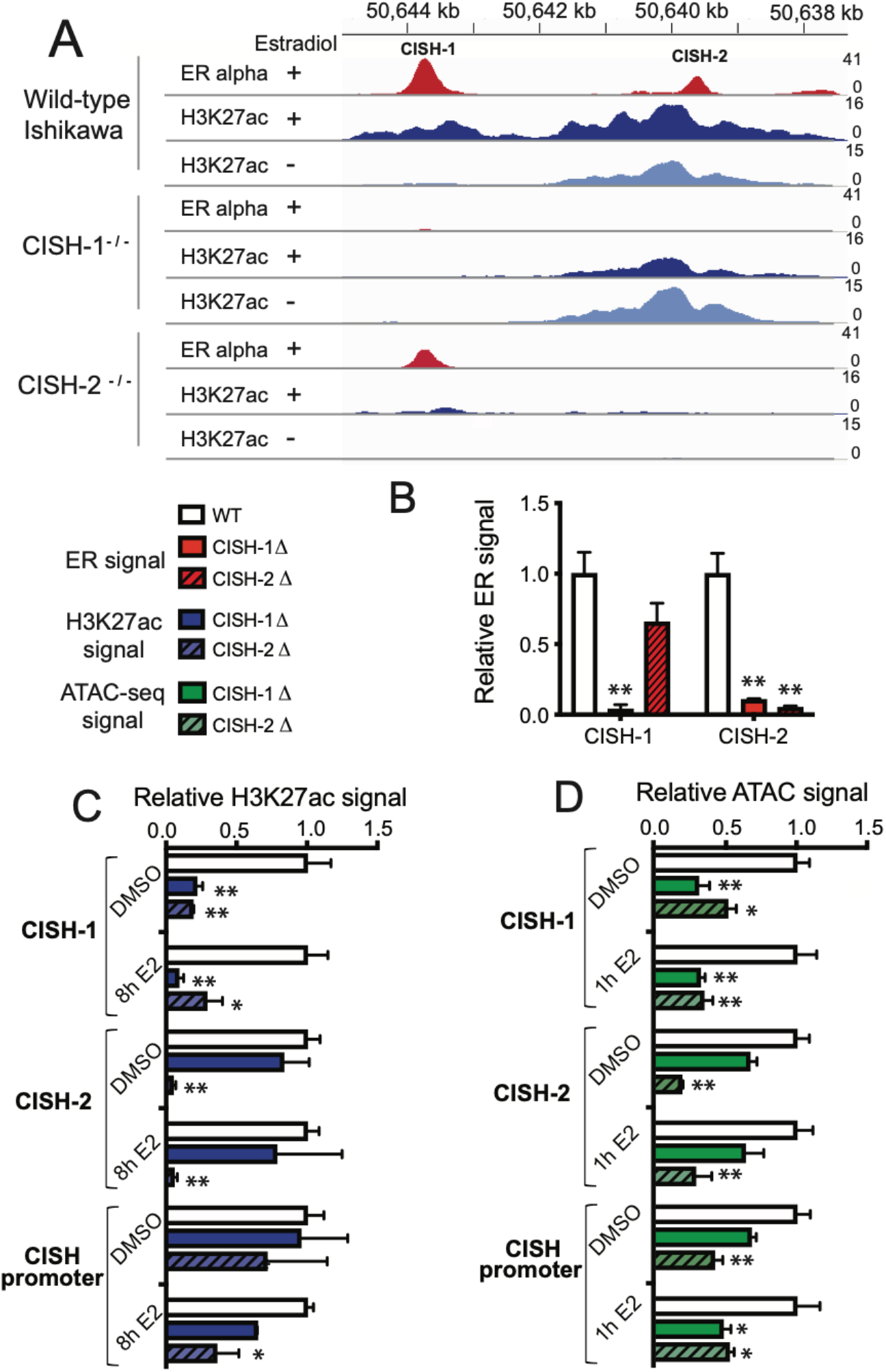
Synergistic enhancers divide tasks of recruiting ER and maintaining permissive chromatin. **A)** Example of the effects of individual *CISH* enhancer deletions on ER binding (red) and H3K27ac (blue) at the neighboring enhancer as measured by ChIP-seq in a representative clone. **B)** ER ChIP-seq signal from 2 independent cell lines containing each deletion (solid red for CISH-1 and striped for CISH-2) was normalized to wild-type cells (white). **C)** H3K27ac ChIP-seq signal from 2 independent cell lines containing each deletion (solid blue for CISH-1 and striped for CISH-2) and treated with either DMSO or 10 nM E2 for 8h was normalized to wild-type cells (white). **D)** ATAC-seq signal from 2 independent cell lines containing each deletion (solid green for CISH-1 and striped for CISH-2) and treated with either DMSO or 10 nM E2 for 1h was normalized to wild-type cells (white). In B-D, error bars represent SEM, * indicates p < 0.05, and ** p < 0.001 in a two-way ANOVA.

We next investigated how enhancer CISH-2 contributes to the synergistic relationship at *CISH*. Deleting 166 bp of CISH-2 resulted in loss of ER binding, H3K27ac and chromatin accessibility at CISH-2 as expected (Figure 3A-D). Loss of CISH-2 caused ER signal at CISH-1 to be reduced by 34% (p-value > 0.05) with ER binding still detectable (Figure 3B). Deletion of CISH-2 led to loss of H3K27ac signal at CISH-1, with or without E2 induction, to levels seen upon CISH-1 loss, indicating that CISH-2 is necessary for active chromatin at CISH-1 (Figure 3C). In addition, CISH-2 loss had a greater effect on *CISH* promoter H3K27ac levels than CISH-1 loss with or without E2 treatment (Figure 3C), suggesting that CISH-2 may be responsible for H3K27ac levels at the entire locus. Accessibility of CISH-1 was also significantly affected by CISH-2 loss, with a 50% loss in the absence of estrogen and a 66% loss in the presence of estrogen (Figure 3D). CISH-2 deletion had a greater impact on promoter accessibility prior to E2 treatment; however, CISH-1 and CISH-2 were equally necessary for promoter accessibility in the presence of estrogen, with a 50% reduction in accessibility in the absence of either site (Figure 3D). CISH-2 exhibits open and acetylated chromatin prior to E2 treatment, while CISH-1 does not, indicating that these features may be important for CISH-2 function. These results suggest that the synergy between CISH-1 and CISH-2 is mediated by the sharing of ER recruitment and the maintenance of active and accessible chromatin between enhancers. CISH-1 contributes ER recruitment, which is consistent with the presence of a full ERE, while CISH-2 promotes a permissive chromatin environment. Altogether, the results at *CISH* indicate that recruiting ER to the genome is not sufficient for an ERBS to produce the estrogen response: some level of histone acetylation and accessibility, here provided by CISH-2, is required for a site to contribute to gene regulation.

### ER binding and active chromatin at supportive enhancer MMP17-2 require predominant enhancer MMP17-1

We next explored the molecular underpinnings of the hierarchical relationship between enhancers at *MMP17*. Loss of 150 bp of MMP17-1 abolished ER binding and greatly reduced H3K27ac and ATAC-seq signal after estrogen treatment at MMP17-1 as expected (Figure 4A-D). MMP17-1 deletion reduced ER binding at MMP17- 2 by 79% compared to wild-type levels (Figure 4A-B). MMP17-1 deletion also greatly reduced H3K27ac levels at MMP17-2, leading to a 64% reduction in the absence of E2 and a 78% reduction in the presence of E2 (Figure 4C). These results suggest that MMP17-1 controls both active chromatin and ER recruitment at MMP17-2. Chromatin accessibility in the absence of E2 was not significantly affected by MMP17-1 loss, but the accessibility of both MMP17-1 and MMP17-2 was reduced by half after E2 treatment (Figure 4D). At the promoter of *MMP17*, histone acetylation levels were reduced by 85% upon MMP17-1 loss in the context of E2 treatment, while promoter accessibility was not significantly affected in either condition, suggesting that other factors are likely involved in maintaining open chromatin at the promoter. Together these results demonstrate that MMP17-1 is at the top of the hierarchy because it orchestrates ER recruitment (through a full ERE), histone acetylation, and, to a lesser extent, chromatin accessibility of MMP17-2, explaining why MMP17-1 is needed in order for MMP17- 2 to contribute to the estrogen response.

**Figure 4.**
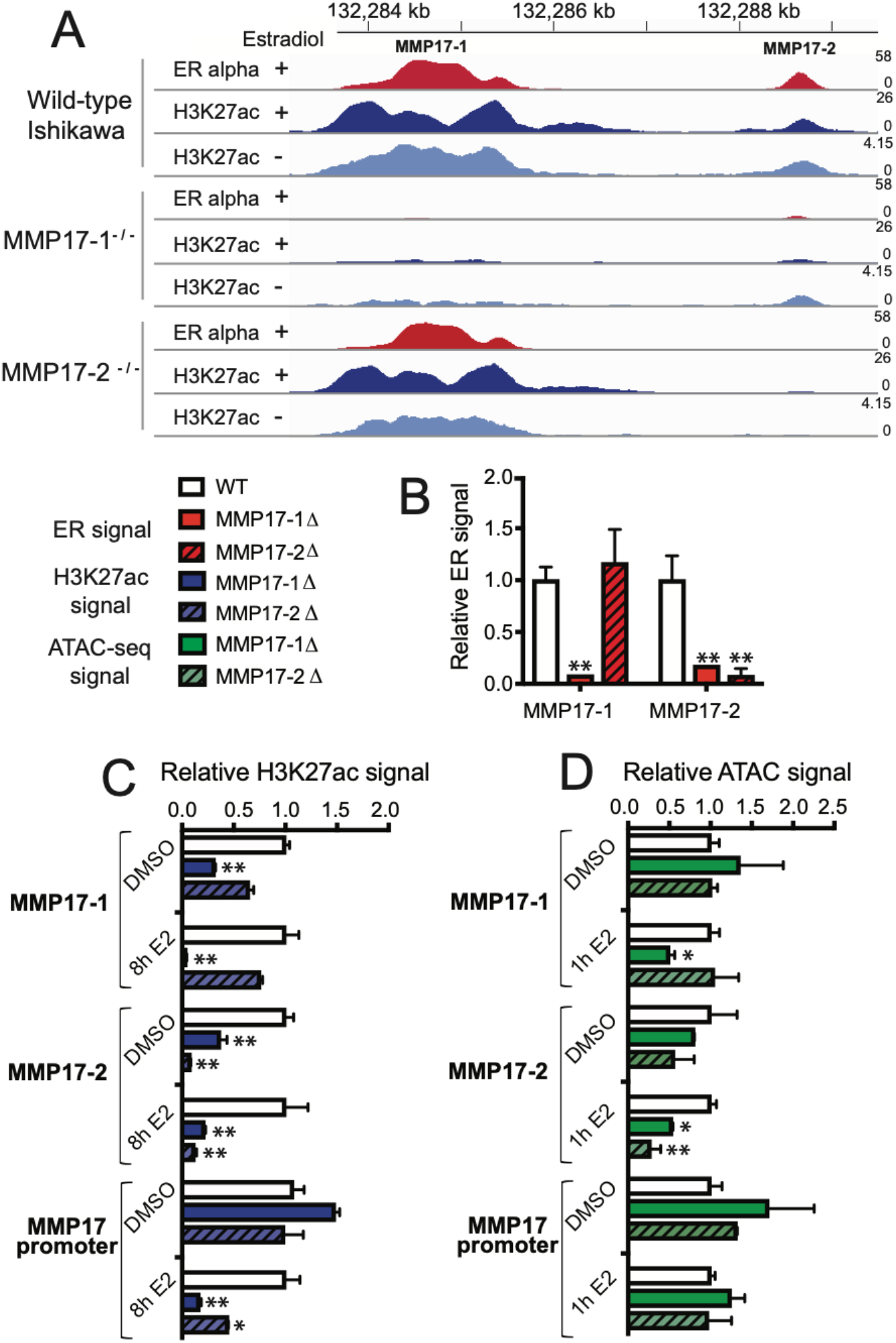
Predominant ERBS regulating *MMP17* controls ER binding and activity at the neighboring site. **A)** Example of the effects of individual *MMP17* enhancer deletions on ER binding (red) and H3K27ac (blue) at the neighboring enhancer as measured by ChIP-seq in a representative clone. **B)** ER ChIP-seq signal from 2 independent cell lines containing each deletion (solid red for MMP17-1 and striped for MMP17-2) was normalized to wild-type cells (white). **C)** H3K27ac ChIP-seq signal from 2 independent cell lines containing each deletion (solid blue for MMP17-1 and striped for MMP17-2) and treated with either DMSO or 10 nM E2 for 8h was normalized to wild-type cells (white). **D)** ATAC-seq signal from 2 independent cell lines containing each deletion (solid green for MMP17-1 and striped for MMP17-2) and treated with either DMSO or 10 nM E2 for 1h was normalized to wild-type cells (white). In B-D, error bars represent SEM, * indicates p < 0.05, and ** p < 0.001 in a two-way ANOVA.

We next examined how MMP17-2 might be acting as a supportive site at the locus. We found that loss of 128 bp of MMP17-2 did not significantly affect ER binding at MMP17-1, indicating that it is not required for the initial recruitment of ER to the locus (Figure 4A-B). MMP17-2 deletion resulted in a 35% reduction of H3K27ac at MMP17-1 in the absence of estrogen (Figure 4C), suggesting that MMP17-2 loss affects MMP17-1 to some extent. However, ER binding at MMP17-1 appears to be able to partially compensate for the reduced histone acetylation, as MMP17-2 loss does not significantly reduce H3K27ac levels at MMP17-1 after E2 treatment. Deletion of MMP17-2 led to 55% reduction of H3K27ac at the *MMP17* promoter in the presence of E2, indicating that loss of MMP17-2 does affect the promoter; however, chromatin accessibility is not significantly impacted by MMP17-2 loss, similar to MMP17-1 loss (Figure 4D). This data leads to a model in which MMP17-1 is responsible for ER recruitment and active chromatin in the neighborhood. MMP17-1 is able to confer these features to MMP17-2, allowing MMP17-2 to contribute to gene regulation as well.

### DNA sequence is critical in determining how an ER-bound enhancer contributes to gene expression

The analysis of enhancer deletions uncovered a molecular interplay between enhancers with ERBS contributing ER recruitment and/or active and accessible chromatin to their neighborhood. We next sought to determine the ERBS features that govern each site’s contribution to gene expression, considering two possibilities: 1) The underlying DNA sequence is dictating the contribution of a site, or 2) the genomic location of a site determines the site’s role in gene regulation. To investigate the importance of these features, we used Cas9 to edit the sequence of ERBS at *MMP17* and *CISH* to match the core ~200 basepairs of the neighboring enhancer. We used Knock-in Blunt Ligation (Geisinger et al., 2016), which relies on double-stranded break repair by non-homologous end joining to insert a PCR product into the desired locus. While the orientation of the insert cannot be controlled this way, ER binds palindromic sequences and enhancers can work in either orientation, suggesting that orientation should not have a large impact on gene regulation.

We created two independent cell lines where one allele of CISH-1 had been replaced with the core sequence of CISH-2. For both of these cell lines, the other allele of CISH-1 was deleted. We compared the estrogen response of these lines to wild-type lines as well as clones containing deletions of CISH-1. We found that the insertion of CISH-2’s sequence did not create an estrogen response in the absence of CISH-1’s core sequence in either clone (Figure 5A), indicating that the sequence of CISH-1 is required for an estrogen response. Similarly, at *MMP17*, replacing one MMP17-1 allele with the core sequence of MMP17-2 led to a loss of the estrogen response when the other MMP17-1 allele was deleted (Figure 5B). Even when one MMP17-1 allele was present with MMP17-2 on the other allele, the response was reduced by 75% compared to wild-type levels, indicating that a fully wild-type MMP17-1 sequence is needed for the estrogen transcriptional response. These results are consistent with our analysis of enhancer deletions, where the full EREs present in MMP17-1 and CISH-1 are required for ER recruitment to the neighborhood and half EREs cannot compensate for the lack of a full ERE.

**Figure 5.**
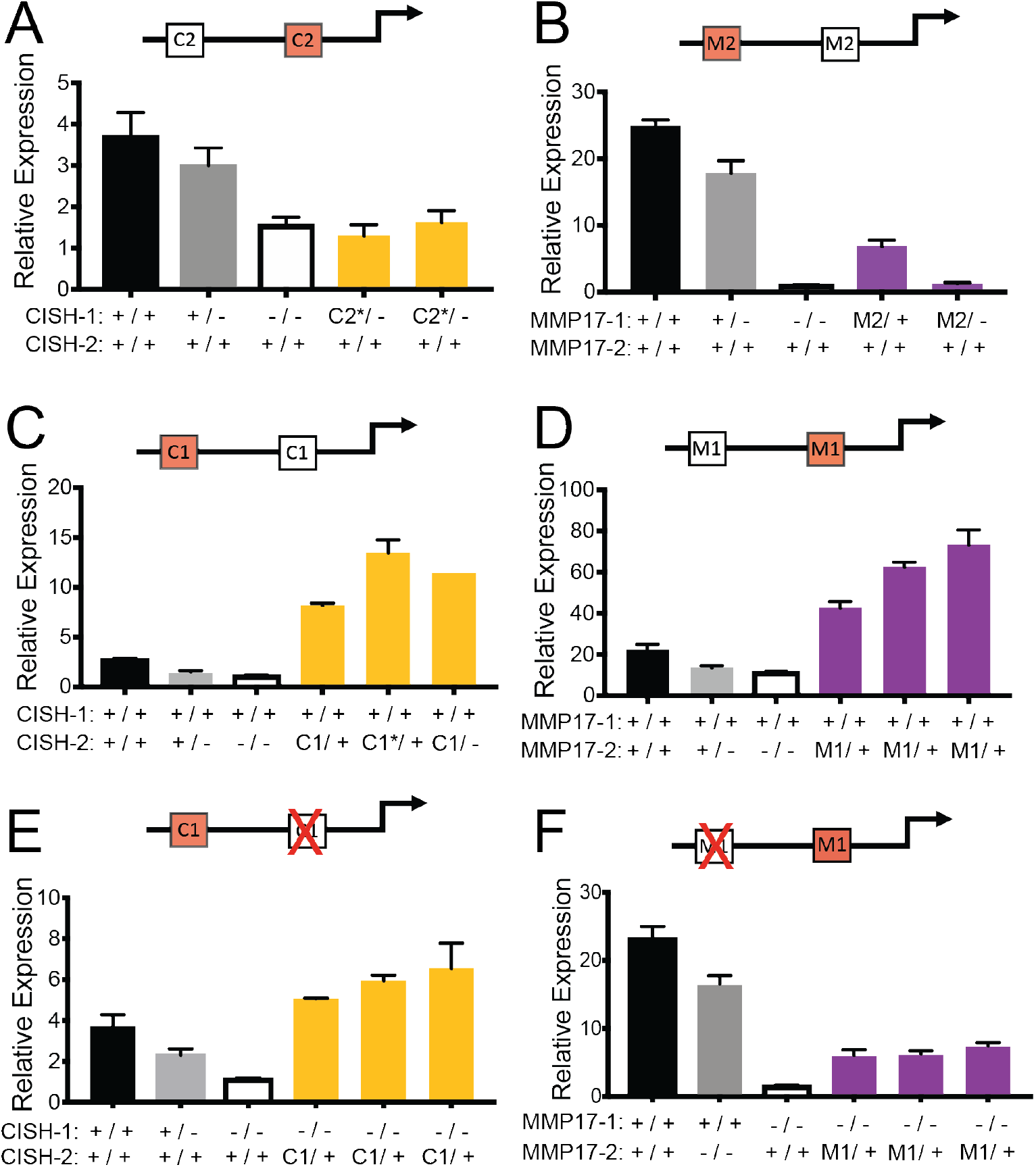
ERBS sequence and location are both important in driving the transcriptional response to estrogen. Red sites on the diagrams have been knocked-in and a red “X” represents deletion. **A – D)** The impact on the transcriptional response to E2 of replacing one allele of an enhancer with the sequence of the neighboring enhancer. This includes replacing CISH-1 with the sequence of CISH-2 (A, yellow bars), MMP17-1 with the sequence of MMP17-2 (B, purple bars), CISH-2 with the sequence of CISH-1 (C, yellow bars), and MMP17-2 with the sequence of MMP17-1 (D, purple bars). Wild-type (black), heterozygous deletion (gray), and homozygous deletion (white) clones are shown for each gene. **E)** The gene regulatory impact of replacing one allele of CISH-2 with the sequence of CISH-1 (yellow bars) in the absence of wild-type CISH-1 is shown for multiple independent clones as well as wild-type (black), heterozygous CISH-1 deletion (gray), and homozygous CISH-1 deletion clones (white). **F)** The gene regulatory impact of replacing the sequence of MMP17-2 with MMP17-1 when wild-type MMP17-1 is deleted is shown for multiple independent clones as well as wild-type (black), MMP17-2 deletion (gray), and MMP17-1 deletion (white) clones. In all panels, bars represent qPCR data from 2 biological replicates following an 8h estrogen induction and expression is relative to an 8h vehicle treatment for each clone. * indicates opposite orientation of insertion in terms of native locus relative to the target gene.

We next examined the effects of replacing CISH-2, which maintains active and accessible chromatin in the neighborhood, with the core sequence of CISH-1. When one allele of CISH-2 was replaced with the core sequence of CISH-1, the estrogen response more than doubled relative to wild-type levels in all three clones we isolated (Figure 5C). Similarly, replacing MMP17-2 with the core sequence of MMP17-1 resulted in a doubling or more in expression levels of *MMP17* compared to the wild-type response for all three clones (Figure 5D). These results suggest that the wild-type expression of *MMP17* and *CISH* is suboptimal in terms of levels, as greater expression levels can be achieved when a sequence with a second full ERE is added to a neighborhood. These results also highlight the importance of local DNA sequence in determining ERBS function.

### Genomic location influences the activity of ER-bound enhancers

While DNA sequence is clearly important in determining an ERBS’s contribution to gene regulation, the genomic location of that sequence could also play a large role in the ability of an ERBS to regulate gene expression. To uncover the importance of location in ERBS activity, we next deleted the neighboring wild-type site in the DNA sequence swapped cell lines described above. This approach created alleles in which the core sequence of an ER-bound enhancer had been moved to the neighboring ERBS location without the influence of another ERBS. These manipulations allowed us to specifically examine the effect of a new location for a given ERBS sequence. While CISH-1 in its native location is unable to support an estrogen response when CISH-2 is deleted, moving the core sequence of CISH-1 into the CISH-2 location was sufficient to drive an estrogen response that is greater than wild-type levels, even without another ERBS in the neighborhood (Figure 5E). The ability of a single copy of CISH-1’s core sequence to drive higher expression in CISH-2’s location than in its native location indicates that while CISH-1 may be an optimal sequence for inducing an E2 transcriptional response, its location is suboptimal, potentially due to the lack of H3K27ac at CISH-1 prior to an estrogen treatment.

When we previously deleted MMP17-2, we found that the predominant site MMP17-1 can drive approximately half of *MMP17*’s response when acting alone. However, when MMP17-1’s core sequence is moved from its original location to MMP17-2’s location, this sequence was unable to drive expression at levels achieved by wild-type MMP17-1 without the support of MMP17-2 (Figure 5F). While the clones with the MMP17-1 location change could still contribute to the E2 response to some extent, the level is significantly less than half of MMP17-2 homozygous deletion clones (p-value = 0.0004, *t*-test). These results indicate that MMP17-1 is an optimal location for the underlying optimal sequence of MMP17-1 and that MMP17-2’s location is suboptimal. Interestingly, both optimal locations (MMP17-1 and CISH-2) are in open chromatin surrounded by high levels of H3K27ac prior to estrogen induction, suggesting that a combination of active chromatin and DNA sequence (with a full ERE) combine to drive an estrogen response.

## Discussion

The ER-bound enhancers that we have previously studied in detail are within 5 kb of one another and combine cooperatively to regulate gene expression in response to estrogen. In this study we find that ER-bound sites in endometrial cancer cells tend to cluster together in the genome much more than expected by chance, as previously reported in breast cancer cells (Bojcsuk et al., 2017; Saravanan et al., 2020), suggesting that enhancer collaboration between neighboring ERBS could be common. In order to determine how ER-bound enhancers are impacting one another, we analyzed cell lines with homozygous deletions of two pairs of neighboring enhancers that regulate *CISH* and *MMP17*. We discovered molecular interplay between neighboring enhancers, which provides an explanation for the collaboration that we see between ER-bound enhancers.

One way in which these enhancers impact their neighbors is through recruitment of ER, despite kilobase-pairs of sequence in between them. Such cooperativity is common when binding motifs are immediately adjacent to one another. For example, deletion of a single binding site for OCT4/SOX2 within an enhancer near the murine *Klf4* locus prevents other transcription factors from binding to that same enhancer and reduces chromatin accessibility at the enhancer (Xie et al., 2017). Enhancers can also impact each other at larger distances, such as *PU.1* autoregulation in myeloid cells, where C/EBPa binds to an enhancer 14 kb upstream of *PU.1* and promotes the activation and accessibility of a neighboring enhancer 2 kb downstream of the predominant enhancer (Leddin et al., 2011). Similar enhancer hierarchies between individual enhancers have recently been described in other systems (Huang et al., 2018; Iwata et al., 2017), and our genome-wide observations suggest that these hierarchies are likely a consistent feature of estrogen-dependent gene regulation.

For both *MMP17* and *CISH*, the enhancer required for bringing ER to the ERBS cluster contained a full ERE, while the ERBS with a half ERE was dispensable for ER binding. When the DNA sequences of full ERE sites were replaced with half ERE containing sequences, the gene expression response to estrogen was lost. Alternatively, swapping a half ERE containing core sequence with a full ERE containing sequence resulted in a super-response to estrogen for both genes. While this indicates that the enhancer sequences of *MMP17* and *CISH* are suboptimal in terms of levels, the combination of sequences allows for fine tuning of an estrogen response through evolutionary changes in both sequence and ERE location. The importance of a full ERE in a cluster is consistent with findings in mice expressing ER DNA binding domain mutants, where only consensus EREs with 1-2 mismatches contribute to gene expression (Coons et al., 2017). In addition, an analysis of ER bound “super enhancers” found that ERE motif strength is higher in regions that precede “super enhancer” formation (Bojcsuk et al., 2017). Our findings indicate that full EREs are necessary for ER recruitment and gene expression, although we also found a role for sites with half EREs. Since our intensive study was limited to two genes, it is possible that multiple half ERE ERBS can combine to regulate expression. Previous studies have found that multiple suboptimal TF binding motifs can drive expression of reporters in *Ciona* and *Drosophila* (Crocker et al., 2015; Farley et al., 2016). However, multiple weak TF binding motifs have yet to be manipulated in the genome simultaneously, and their ability to contribute in enhancer pairs remains unclear.

In addition to neighboring ERBS contributing to each other’s ability to bind ER, ER-bound enhancers can also impact one another’s chromatin environment. CISH-2 and MMP17-1 were needed to maintain histone acetylation and chromatin accessibility in their respective neighborhoods. Clustered ERBS can aid each other in maintaining permissible chromatin and this chromatin environment appears crucial. When core sequences of full ERE enhancers were moved to different locations, we found that placing a full ERE sequence in a region with high basal histone acetylation had a larger impact on transcription than adjacent regions with lower histone acetylation. Our results suggest that sites contributing to the transcriptional response to estrogen need some level of permissible chromatin prior to an estrogen induction, whether it is at the same site as the full ERE (e.g. MMP17-1) or the neighboring site (e.g. CISH-2). These findings are consistent with previous work showing that recruitment of p300 and subsequent histone acetylation appears to be a crucial step in estrogen-dependent gene regulation, as inhibitors of the p300 catalytic domain abrogate estrogen-dependent transcription (Murakami et al., 2017). The fact that permissible chromatin in the absence of estrogens appears important suggests that these sites may be somehow primed for activation; however, it remains unclear what factors are involved in priming these sites. Analysis of additional factors bound to these regions (Figure 2C-D) indicates that p300 is bound to CISH-2 and MMP17-1, the sites responsible for acetylation of the neighborhood. NFIC, TCF12, and ETV4 (Rodriguez et al., 2020) were also uniquely found at CISH-2 and MMP17-1 as opposed to CISH-1 and MMP17-2. Although with only two sets of ERBS, it is unclear if these binding patterns are commonly observed.

When taken together, our observations provide an explanation for the synergistic relationship of ERBS regulating *CISH* and the hierarchical relationship of enhancers regulating *MMP17* (Figure 6). MMP17-1 is responsible for both ER recruitment and an active chromatin environment, which is consistent with its role atop the hierarchy. The activity of MMP17-1 allows MMP17-2 to bind ER and adopt an active chromatin context. CISH-1 and CISH-2 exhibit balanced regulatory sharing, which explains the complete synergy observed in their activation of *CISH* expression. CISH-1 contributes ER recruitment while CISH-2 provides permissible chromatin and both molecular actions are required for the estrogenic transcriptional response of *CISH*. The molecular mechanisms of how both sets of enhancers work together in *cis* reconciles previously disparate observations concerning these ER-bound enhancers: When ER is driving the transcriptional response, enhancers work cooperatively (Carleton et al., 2017) and they do so by contributing to each other’s activity; however, when activating cofactors are directly recruited to these sites, synergy is lost (Ginley-Hidinger et al., 2019) because the cooperative steps (ER recruitment, chromatin accessibility, and histone acetylation) have been bypassed.

**Figure 6.**
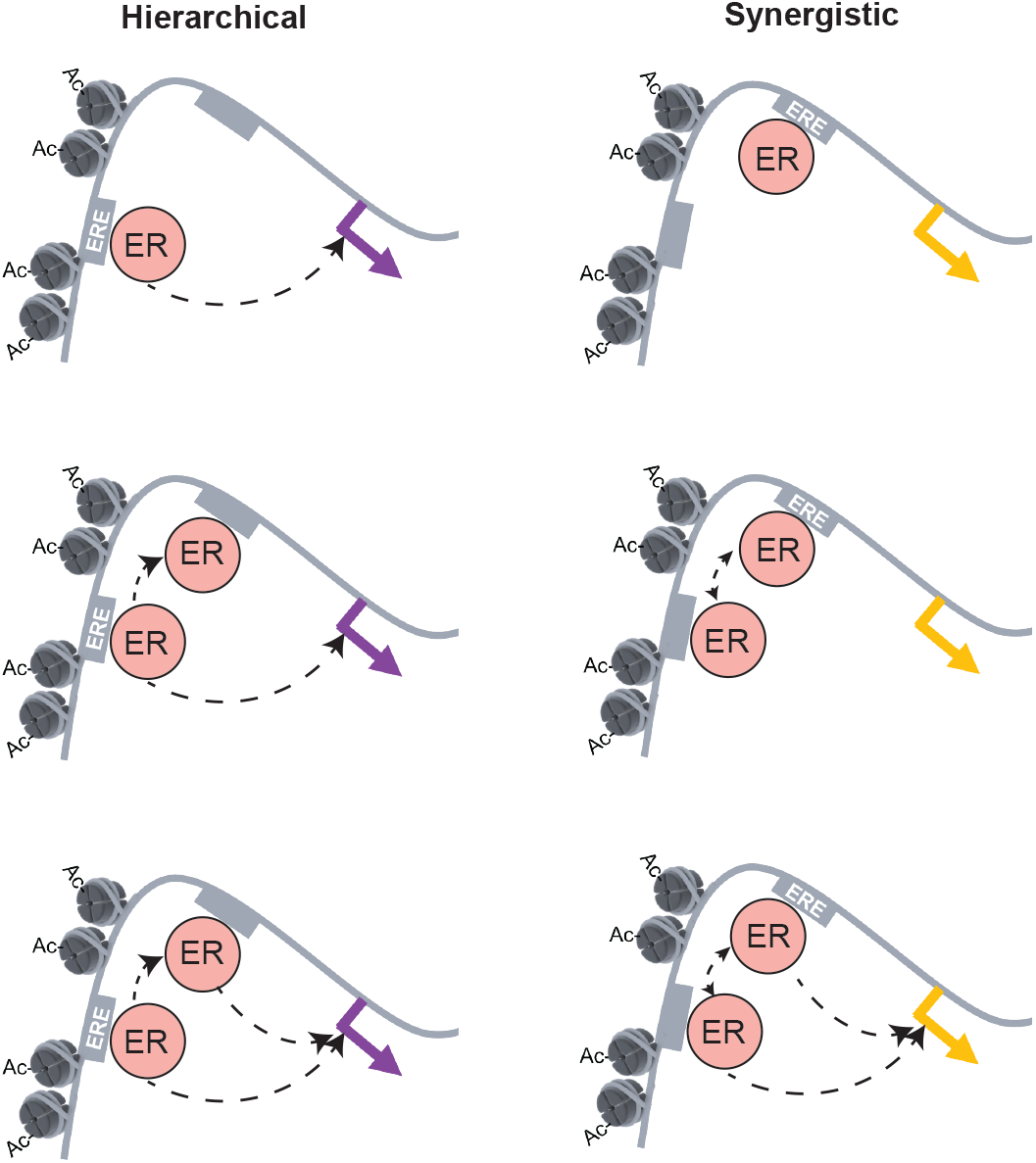
Model for two distinct modes of enhancer collaboration at neighboring ERBS. In a hierarchical relationship, a site in permissive chromatin and containing a consensus ERE motif can directly recruit ER and regulate gene expression. Only after this site becomes active can ER bind at the neighboring site that contains a weak ERE. This site contributes to gene expression by affecting activity of the promoter. In a synergistic interaction, the strong ERE site can recruit ER, but cannot contribute to expression without the presence of the neighboring weak ERE site. The presence of the weak ERE site promotes active chromatin in the region while also acting as a second binding site for ER, allowing both sites to contribute to gene expression.

## Author Contributions

Conceptualization, J.B.C. and J.G.; Methodology, J.B.C. and J.G.; Investigation, J.B.C., M.G-H, K.B., and J.G.; Formal Analysis, J.B.C., M.G-H, R.M.L., and J.G.; Writing – Original Draft, J.B.C. and J.G.; Writing – Review & Editing, all authors.; Supervision, A.R.Q. and J.G.; Funding Acquisition, A.R.Q. and J.G..

## Acknowledgements

This work was supported by NIH/NHGRI R01 HG008974 to J.G. and A.R.Q., and the Huntsman Cancer Institute. Research reported in this publication utilized the High-Throughput Genomics Shared Resource at the University of Utah and was supported by NIH/NCI award P30 CA042014. J.B.C. was supported by NIH/NIGMS Training Program in Genetics T32 GM007464. We thank John Lis and Erin Wissink for their suggestions on the study and the manuscript.

## Methods

### Cell culture

Ishikawa cell lines containing deletions of individual ER binding sites were generated as described previously (Carleton et al., 2017). Ishikawa cells were maintained in RPMI 1640 (Gibco) supplemented with 10% FBS (Gibco) and 1% pencillin/streptomycin (Gibco). Cells were placed in hormone-depleted RPMI (phenol red-free RPMI-1640 supplemented with 10% Charcoal/Dextran treated fetal bovine serum (HyClone) and 1% penicillin-streptomycin) for at least 6 days prior to E2 treatment.

### ChIP-seq

Approximately 20 million cells were plated in 15 cm dishes the day before harvest for each cell line of interest. For ER ChIP-seq, a 1 h 10 nM E2 induction was performed prior to harvest. For H3K27ac ChIP-seq, an 8 h 10 nM E2 or DMSO treatment was performed prior to harvest. To crosslink chromatin, 37% formaldehyde (Sigma-Aldrich) was added directly to the media for a final concentration of 1% and allowed to incubate at room temperature for 10 minutes. Glycine was added at a final concentration of 125 mM to stop crosslinking, and cells were washed with cold 1X PBS. Plates were scraped in Farnham lysis buffer with 1X protease inhibitor (Thermo Fisher). Chromatin was sonicated using an Active Motif EpiShear Probe Sonicator with 6 cycles of 30 s, at 40% amplitude, with 30 s of rest. ChIP was performed as previously described (Reddy et al., 2009) using anti-ER (Santa Cruz HC-20) and anti-H3K27ac (Active Motif 39133) antibodies. Libraries were sequenced on an Illumina HiSeq 2500 as single-end 50 basepair reads. Reads were aligned to hg19 using bowtie (Langmead et al., 2009) with the following parameters: -m 1 -t ‒best -q -S -l 32 ‒e 80 -n 2. MACS2 (Zhang et al., 2008) was used to call peaks with a p-value cutoff of 1e-10 and the mfold parameter bounded between 5 and 50. The input control used in peak-calling was derived from the parental Ishikawa line. The broad peak calling feature was used for H3K27ac ChIP-seq. Bedtools (Quinlan and Hall, 2010) was used to count reads in peaks. A window of 2 kb centered on the summit was used for counting H3K27ac ChIP-seq reads within peaks and a window of 500 bp was used for ER ChIP-seq read counting. ChIP-seq signals of an enhancer deletion was compared to clones that were wildtype for that locus but harbored enhancer deletions nearby the other gene. For example, MMP17-1 enhancer deletions were compared to CISH-1 and CISH-2 deletions as controls. Additional ChIP-seq data (Figure 2C-D) was from the ENCODE project (Consortium, 2012).

### ATAC-seq

For each of the clones, approximately 250,000 cells were lysed and nuclei were harvested for transposition as described previously (Buenrostro et al., 2013) following a 1 h treatment with E2 or DMSO. Tn5 transposase with Illumina adapters was assembled as previously described (Picelli et al., 2014). Libraries were sequenced on a HiSeq 2500 with single-end 50 basepair reads. Reads were aligned to hg19 using bowtie (Langmead et al., 2009) and the following parameters: -m 1 -t–best -q -S -l 32 -e 80 -n 2. SAM files were converted to BAM files for peak calling using samtools (Li et al., 2009). MACS2 (Zhang et al., 2008) was used to call peaks without an input using the following command: macs2 callpeak --nomodel --shift -100 --extsize 200 ‒B ‒SPMR. Peaks were overlapped with ER binding sites and reads were counted using bedtools (Quinlan and Hall, 2010). ATAC-seq signals of an enhancer deletion was compared to clones that were wildtype for that locus but harbored enhancer deletions nearby the other gene.

### Knock in blunt ligation

To generate PCR products for knock-ins of entire ERBS, we designed PCR primers (Table S1) to specifically amplify the 125-225 bp region for each site that was deleted using Cas9 with previously designed guide RNAs (Carleton et al., 2017). The PCR primers (IDT) included phosphothiorate bonds between the first 3 nucleotides to prevent degradation (Geisinger et al., 2016). PCR products were purified using Ampure XP (Beckman Coulter) and Sanger sequenced (Genewiz) to verify the product.

To generate knock-ins at a given ERBS, Ishikawa cells were plated at a density of ~300,000 cells per well in 6 well plates and transfected the following day with 1650 ng Cas9 plasmid (Addgene 62988, a gift from Feng Zhang), 550 ng of each guide RNA (Table S2), and 200 ng PCR product using FuGENE HD (Promega). At 2 days post transfection, the media was changed and supplemented with 1 μg/mL puromycin to select for transfected cells. After 1-2 days of selection, the media was again changed to allow cells to recover for at least 1 day prior to limiting dilution cloning. When cells became confluent, they were subjected to limited dilution cloning to isolate individual colonies containing specific mutations. When colonies were sufficiently large (about 2 weeks following plating), approximately 96 colonies were picked for each knock-in experiment. Colonies were allowed to grow in a 24-well plate until confluent, at which point they were moved to a 48-well plate, and most of the cells were harvested for genomic DNA isolation. Genomic DNA was isolated using the ZR-96 Quick-DNA extraction kit (Zymo Research) and subject to PCR using primers outside of the regions of interest that contained Illumina tails for high-throughput sequencing (Table S1). PCR products were cleaned up using ZR-96 DNA Clean and Concentrate kits (Zymo Research) and a second PCR was performed to attach barcodes. PCR products were pooled by region of interest using 2-3 μL of each reaction and the resulting pools of products were purified using a 1X Ampure XP cleanup (Beckman Coulter). Purified libraries were pooled and submitted for paired-end 150 cycle sequencing on an Illumina MiSeq. Reads were aligned to a custom library containing the inserts of interest using bwa (Li and Durbin, 2010) with the following parameters: bwa mem ‒M ‒t 2.

### RNA isolation and qPCR

To harvest RNA, we used a direct on-plate lysis of cells with 300 μL of Buffer RLT Plus (Qiagen) supplemented with 1% beta-mercaptoethanol (Sigma-Aldrich). Lysates were purified using the ZR-96-well Quick-RNA kit (Zymo Research). RNA was quantified using RiboGreen (ThermoFisher Scientific) on a Wallac EnVision plate reader (PerkinElmer) or on a Qubit 2.0 (ThermoFisher Scientific). Gene expression was quantified using Power SYBR Green RNA-to-CT 1-Step Kit (ThermoFisher Scientific) and a CFX Connect light cycler (BioRad). Each reaction contained 50 ng of RNA as starting material. As per kit instructions, 40 cycles of PCR were performed following a 30-minute cDNA synthesis. Primers for *CTCF*, *CISH*, and *MMP17* can be found in Table S1.

### Analysis of ERBS clusters and their activity

To identify distances between ERBS, we used bedtools closest (Quinlan and Hall, 2010) on previously generated ChIP-seq data for ER following a 1h E2 treatment (Gertz et al., 2013). ERBS clusters were generated using bedtools merge with a distance of 10 kb on a file containing all ERBS. Controls were performed by randomly selecting 8,621 (the number of ERBS) previously identified DNase I hypersensitive sites or CTCF ChIP-seq sites (Gertz et al., 2013) and then performing the same cluster analysis.

### Quantification and statistical analysis

To quantify differences in ChIP-seq and ATAC-seq signal resulting from ERBS manipulations, we used two-way ANOVA. To quantify differences in E2-induced gene expression following genetic manipulation, we used two-way ANOVA. To quantify differences in fold change in response to E2 for upregulated genes with specific numbers of ERBS clusters within 100 kb of their TSSs, we used Wilcoxon rank sum tests. P-values from these tests can be found in the text and in the figure legends.

### Data access

Raw and processed sequencing data can be found at the Gene Expression Omnibus under the following accession numbers: GSE147141 (ChIP-seq) and GSE147140 (ATAC-seq).

**Table S1.**
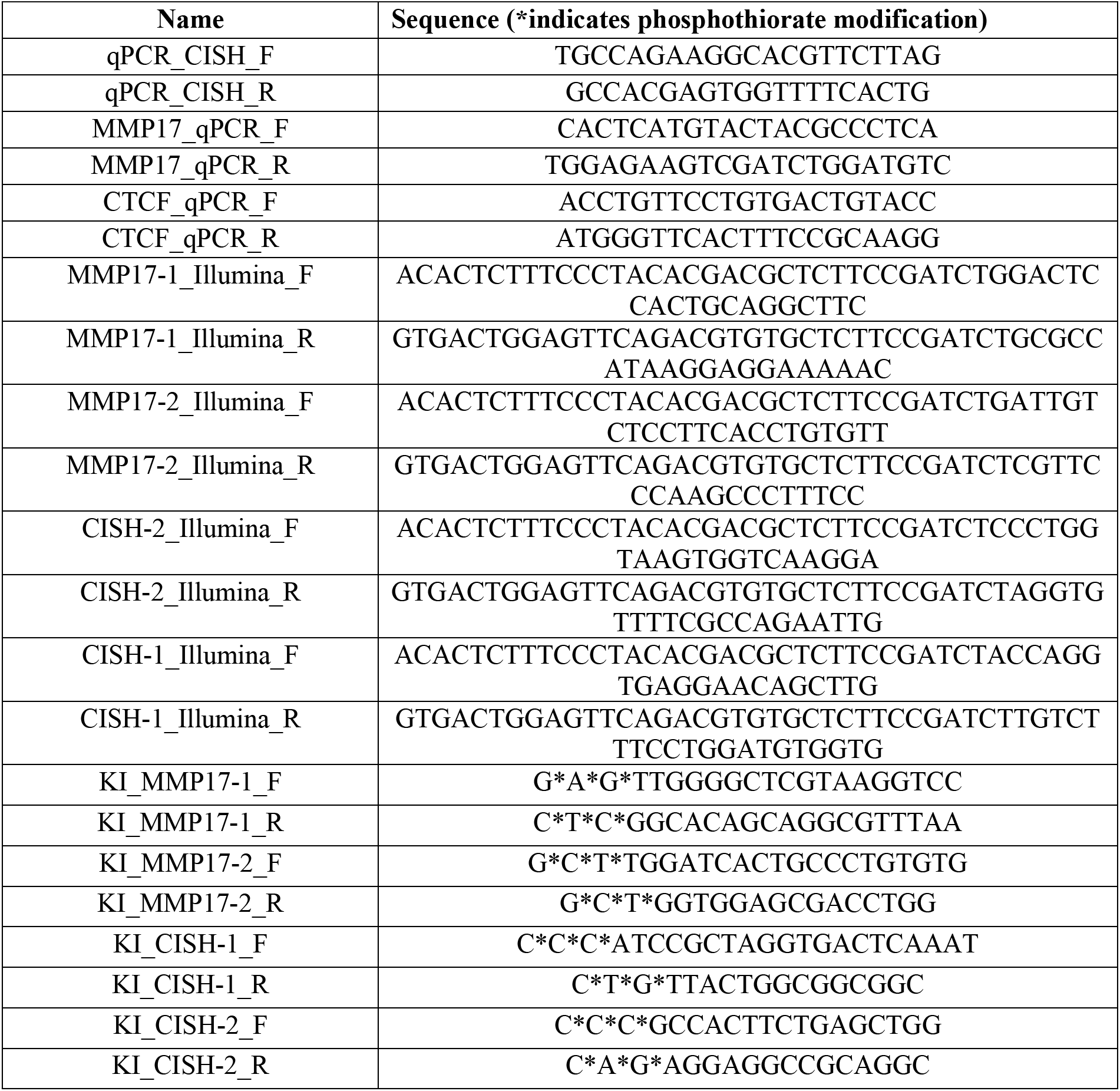
Primers used.

**Table S2.**
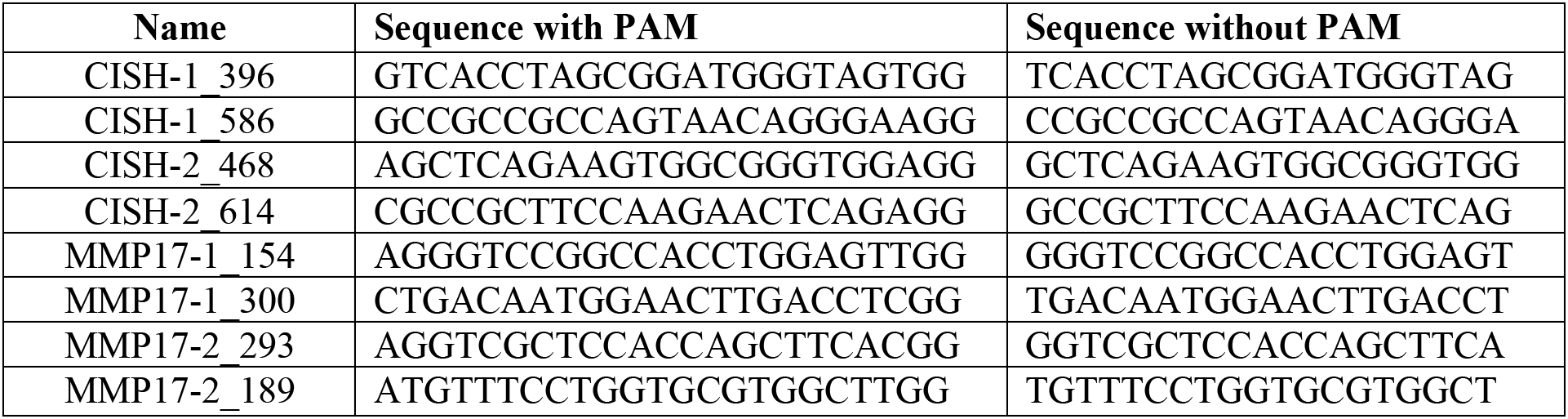
Guide RNAs used.

